# Predicting the conformational flexibility of antibody and T-cell receptor CDRs

**DOI:** 10.1101/2025.03.19.644119

**Authors:** Fabian C. Spoendlin, Monica L. Fernández-Quintero, Sai S. R. Raghavan, Hannah L. Turner, Anant Gharpure, Johannes R. Loeffler, Wing K. Wong, Alexander Bujotzek, Guy Georges, Andrew B. Ward, Charlotte M. Deane

## Abstract

Many proteins are highly flexible and their ability to adapt their shape can be fundamental to their functional properties. We can now computationally predict a single, static protein structure with high accuracy. However, we are not yet able to reliably predict structural flexibility. A major factor limiting such predictions is the scarcity of suitable training data. Here, we focus on predicting the structural flexibility of the functionally important antibody and T-cell receptor CDR3 loops. We extracted a dataset of CDR3 like loop motifs from the PDB to create ALL-conformations, a dataset containing 1.2 million structures and more than 100,000 unique sequences. Using this dataset, we develop ITsFlexible a method classifying CDR3 flexibility, which outperforms all alternative approaches on our crystal structure datasets and successfully generalises to MD simulations. We also used ITsFlexible to predict the flexibility of three completely novel CDRH3 loops and experimentally determined their conformations using cryo-EM.

## 1 Introduction

Many proteins are flexible molecules which adopt several stable structures, termed conformations, and transitions between them can be fundamental for their function (Wei et al., 2016; Teilum et al., 2009). Antibodies and TCRs primarily engage their targets through six loop motifs called the complementarity-determining regions (CDRs) (Chiu et al., 2019). Structural flexibility of CDRs has been linked to several key functional properties. For some antibodies, conformational changes are known to be required for antigen recognition (Liu et al., 2024). The ability of an antigen receptor to adopt multiple conformations has also been associated with polyspecificity as different structural states allow for greater variability in the recognised antigens (Guthmiller et al., 2020; James et al., 2003). Furthermore, flexibility has an effect on binding affinity as it directly impacts the entropic costs of antigen binding (Mikolajek et al., 2022) and rigidification has been observed as one of the natural mechanisms used to increase affinity (Schmidt et al., 2013; Li et al., 2022).

Specificity and affinity are two critical properties of antibody and TCR therapeutics. In order to maximise target and minimise off-target interactions a therapeutic should have high affinity and specificity (Sormanni et al., 2018). This suggests therapeutics should have a preference towards rigidity. However, there is some evidence indicating that conformational flexibility allows better recognition of mutated antigen variants and could be desired when designing broadly neutralising antibodies (Li et al., 2015; Prigent et al., 2018). In either case, a method predicting CDR flexibility would allow both an enhanced investigation of antibody function and the potential to tune desired therapeutic properties.

Computational predictions of a single, static structure of a protein from its sequence is now considered a routine task (e.g. Jumper et al., 2021; Lin et al., 2023; Baek et al., 2021) and more recently developed tools are also showing promise at protein complex prediction (e.g. Abramson et al., 2024; Hayes et al., 2024). However, predicting structures of more than one conformational state remains challenging. One factor which has limited progress in conformation prediction is the scarcity of suitable data.

Evidence on conformational flexibility can currently be obtained from several experimental sources. Nuclear magnetic resonance (NMR) spectroscopy and hydrogen-deuterium exchange mass spectrometry (HDX-MS) can be used to measure protein dynamics in solution although they do not typically provide atomically resolved flexibility information (see Kay (2011) for a review of NMR and Konermann et al. (2011) of HDX methods). X-ray crystallography is the standard method used to obtain high-resolution structures of conformational states. These can be captured by solving separate structures of the same protein under different conditions. Because multiple structures must be available, the number of proteins for which flexibility can be assessed from crystallographic data is much smaller than the total number of solved structures. Crystal structures have been used to explore the flexibility of specific loop types (Marks et al., 2018) or for case studies of full length proteins (Riccabona et al., 2024; del Alamo et al., 2022; Stein & Mchaourab, 2022) but there has been little systematic mining of the protein data bank (PDB) for all instances of the same sequence representing alternative conformations (Ellaway et al., 2023).

Molecular dynamics (MD) simulations provide a computational way to generate conformational ensembles. MD simulations are computationally expensive and, therefore, even the largest databases of standardised MD simulations are not yet sufficient for training machine learning models (Vander Meersche et al., 2024).

Despite these data challenges, a number of methods have been developed that attempt to predict structures of protein conformational ensembles. Recent approaches have concentrated on modifying the AF2 inference procedure to increase the diversity of outputs (e.g. del Alamo et al., 2022; Stein & Mchaourab, 2022; Wayment-Steele et al., 2024; Sala et al., 2023b; Faezov & Dunbrack, 2023; Sala et al., 2023a; Saldaño et al., 2022). These methods target the multiple sequence alignment (MSA), one of the main inputs of AF2, from which co-evolutionary signals are extracted to infer protein residues likely to be in close proximity. In theory, the MSA should contain co-evolutionary information for all conformational states. Deconvolving these signals is generally attempted by reducing the depth of the MSA, for example through random subsampling (del Alamo et al., 2022) or sequence clustering (Wayment-Steele et al., 2024). More recently a range of methods were specifically trained for the task of conformation prediction (e.g. Lewis et al., 2024; Zhu et al., 2024; Zheng et al., 2024; Lu et al., 2024; Huguet et al., 2024). These generally take the form of generative protein structure prediction models trained on the PDB and a small number of molecular dynamics (MD) simulations.

Detailed evaluation of conformation prediction tools is complicated by data scarcity but available evidence suggests that reliable predictions are not yet possible. Many methods have only been evaluated on one or a few case studies which may not accurately reflect their predictions across diverse sets of proteins (Riccabona et al., 2024; Faezov & Dunbrack, 2023; Sala et al., 2023a; Saldaño et al., 2022; Zheng et al., 2024). Evaluation of some methods on slightly larger test sets show that the diversity of predicted structures is generally increased, however the conformational landscape is not covered with high accuracy (Jing et al., 2024). Prediction of conformational states of antibody and TCR CDRs specifically has only been assessed on a handful of case studies (Riccabona et al., 2024).

In this work, we focus on the flexibility of the functionally important antibody and TCR CDR3s. In an attempt to address the issue of data scarcity we consider loop motifs with the same secondary structure pattern, defined as loops bounded by two antiparallel *β*-strands, across all proteins. Through a systematic mining of the PDB (Berman, 2000) and antibody and TCR specific databases (Dunbar et al., 2014; Leem et al., 2018) we created **A**ntibody **L**ike **L**oop conformations (ALL-conformations) a dataset containing 1.2 million crystal structures of loops with 100,000 unique sequences. The dataset captures all experimentally observed conformations of loop motifs found between pairs of antiparallel *β*-strands including both antibody and TCR CDR3s (Figure 1, Panel A). We analysed the structural flexibility in the ALL-conformations set and label more than 20,000 unique loop sequences by their ability to undergo conformational changes (Figure 1, Panel B). Using this data, we built **I**mmunoglobulins and **T**CR**s Flexib**ility c**l**assifi**e**r (ITsFlexible) a method that classifies whether CDR3s are rigid (adopt a single conformation) or exhibit flexibility (transition between multiple states) (Figure 1, Panel C). ITsFlexible predicts the flexibility of CDR3s evaluated in ensembles of crystal structures with high accuracy and achieves state-of-the-art performance. The model also generalises effectively to a test set derived from MD simulations. Furthermore, we used ITsFlexible to predict the flexibility of three completely novel CDRH3 loops and used cryo-EM to experimentally determine their conformations. These experiments showed that two of the three model predictions were correct. ALL-conformations is released on Zenodo (doi.org/10.5281/zenodo.15032263) and ITsFlexible is available on Github (github.com/oxpig/ITsFlexible).

**Figure 1:**
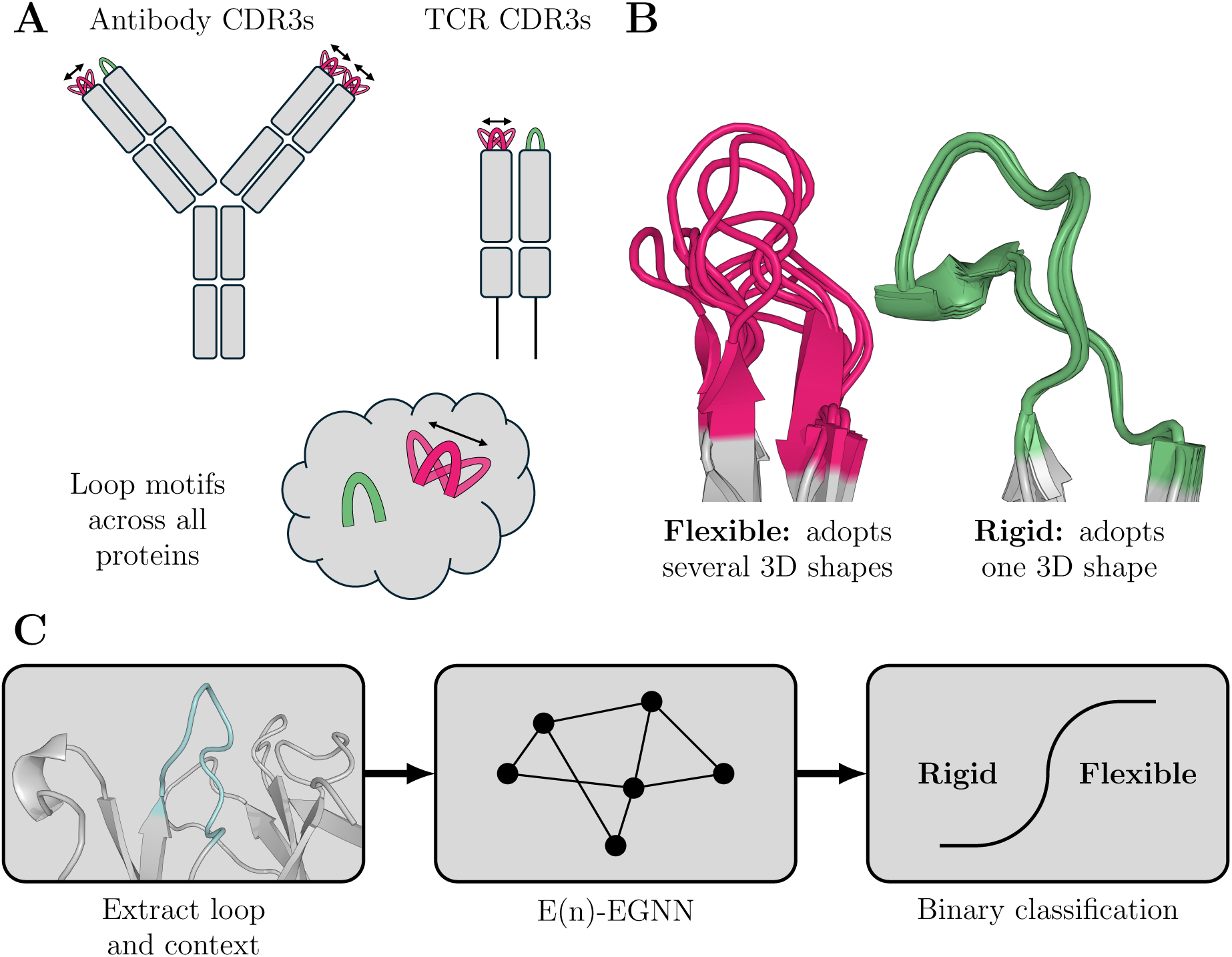
Overview of ALL-conformations and ITsFlexible. A) ALL-conformations is a dataset that contains crystal structures of antibody CDR3s, TCR CDR3s and CDR-like loop motifs across all proteins. The dataset captures all observed conformational states of such loops. B) Loops are labelled as either flexible, if they are observed in more than one conformation, or rigid, if evidence suggest they adopt a single conformation. We define a conformation by structural similarity and use an RMSD of 1.25 Å as a threshold to separate states. C) Flowchart detailing the ITsFlexible method predicting the conformational flexibility of CDR loops. The structure and sequence of a loop (cyan) and its context (grey) are extracted from a PDB file and a graph representation is generated. A graph neural network (GNN) classifies loops as conformationally flexible or rigid.

## 2 Results

### 2.1 ALL-conformations dataset

**A**ntibody **L**ike **L**oop conformations (ALL-conformations) is a dataset that captures the conformational flexibility of loop motifs bounded by two antiparallel *β*-strands which included five subsets, antibody CDRH3s and CDRL3s, TCR CDRB3s and CDRA3s and loop motifs across all proteins in the PDB (Figure S3). The dataset collects available crystal structures of such loops and contains the structures of all experimentally observed conformational states.

Extracting CDR3 structures from SAbDab (Dunbar et al., 2014) and STCRDab (Leem et al., 2018) and loop motifs from the PDB (Berman, 2000) we obtain a total of more than 1.2 million examples with more than 100,000 unique sequences (Table 1). The PDB set contains loops between 1 and 87 residues, however, it is heavily enriched for shorter loops (Figure S4). Length distributions are different for CDR3s with a peak around 10 to 15 residues and no loops shorter than 4 amino acids (Figure S5). We found that a large number of loop sequences are represented by multiple crystal structures, in extreme cases there are several thousand structures for a single loop sequence (Figure S4 & S5).

**Table 1:** Number of structures and sequences in ALL-conformations.

| DATASET | STRUCTURES | UNIQUE LOOP/LOOP + ANCHORS/Fv SEQUENCES | LOOP/LOOP + ANCHORS/Fv SEQS WITH MULTIPLE STRUCTURES |
| --- | --- | --- | --- |
| PDB SET | 1,208,000/- | 99,000/162,000/- | 69,000/112,000/- |
| ANTIBODIES |  |  |  |
| CDRH3 | 6401 | 2190/2190/2477 | 1372/1372/1460 |
| CDRL3 | 6401 | 1762/1776/2366 | 1147/1150/1432 |
| TCRs |  |  |  |
| CDRB3 | 691 | 197/197/234 | 137/137/144 |
| CDRA3 | 691 | 192/196/233 | 134/137/151 |

The loops were classified as rigid, flexible or unknown (see methods for details). We identified more than 4,000 ‘flexible’ loops, those where multiple conformations are observed in crystal structures (Table S5). Given the available data it is not possible to guarantee that all loops labelled to be ‘rigid’ only occupy a single conformation as there always remains a possibility that additional conformational states have not yet been captured by an experimental structure. In our definition, we take only those loops that adopt the same conformation in more than five structures which should ensure that the set is heavily enriched for loops with a single accessible conformation. Using this definition we identify more than 16,000 ‘rigid’ loops (Table S5).

### 2.2 Predicting the flexibility of CDR-like protein loop motifs

The ALL-conformations set was used to train **I**mmunoglobulins and **T**CR**s Flexib**ility c**l**assifi**e**r (ITsFlexible), a model that predicts protein loop flexibility. ITsFlexible is a graph neural network (GNN) (Figure 3) that binary classifies loops if they can occupy multiple accessible conformations (flexible) or adopt a single stable state (rigid) from inputs encoding the sequence and structure of a loop and its structural context (see methods for details).

We trained and evaluated ITsFlexible on the PDB set of ALL-conformations. The data split was performed based on sequence identity and test set loops had a maximum of 80% sequence identity with length matched loops in the training and validation sets. Classifier performance was compared to three baseline models (Table 2) predicting flexibility from biophysical features of length and solvent exposure which have previously been shown to influence loop dynamics (Guloglu & Deane, 2023; Marks et al., 2018). Long loops contain more bonds around which they can rotate and solvent exposure reduces steric hindrance which restricts conformational rearrangements. Therefore, greater length and solvent exposure both tend to promote flexibility (Guloglu & Deane, 2023; Marks et al., 2018). ITsFlexible is able to classify loop flexibility (PR AUC of 0.60 and a ROC AUC of 0.82) and outperforms our baseline classifiers.

**Table 2:** Classification performance on the PDB test set (n = 2845)

| METHOD | ROC AUC | PR AUC |
| --- | --- | --- |
| RANDOM | 0.50 | 0.23 |
| BASELINES |  |  |
| SOLVENT EXPOSURE | 0.60 | 0.29 |
| LENGTH | 0.69 | 0.35 |
| COMBINED | 0.76 | 0.46 |
| ITsFLEXIBLE |  |  |
| ITsFLEXIBLE-SEQUENCE | 0.76 | 0.47 |
| ITsFLEXIBLE-LOOP | 0.76 | 0.46 |
| ITsFLEXIBLE | **0.82** | **0.62** |

The best performance achieved for each metric is highlighted in bold.

We investigated the features predictive of loop flexibility through ablation of model inputs. ITsFlexible-loop has the same architecture as the default model but inputs are reduced to the structure and sequence of only the loop itself (see SI for details). ITsFlexible-sequence is a CNN-based model trained on only a sequence encoding of the loop (see SI for details). ITsFlexible-loop and ITsFlexible-sequence achieve a similar performance to one another. Both outperform the relevant baseline (only baseline-length is relevant here as solvent exposure depends on the structural context context which ITsFlexible-loop and ITsFlexible-sequence do not consider.) but are less predictive than ITsFlexible (Table 2).

These results give some indication on the factors affecting the conformational flexibility of protein loops. It is well known that long loops and loops with greater solvent exposure tend to be more flexible than shorter and more buried ones (Marks et al., 2018; Guloglu & Deane, 2023). While our findings agree with these general trends, we also show that loop sequence is more predictive of flexibility than length alone suggesting the sequence of a loop impacts its ability to adopt multiple conformations. Adding a structural encoding of the loop itself in addition to a sequence encoding does not improve classification. This shows that no information is gained from the loop structure. A large boost in performance was achieved by encoding the structural context of the loop motif. In line with previous evidence from MD studies (Guloglu & Deane, 2023), this highlights that interactions of a loop with its context within the protein are important determinants of its conformational dynamics.

### 2.3 ITsFlexible is highly predictive of CDR3 flexibility

We next investigated the ability of ITsFlexible, trained on general proteins loops in the PDB, to predict the flexibility of antibody and TCR CDR3s. The model was evaluated on the CDR3 sets of ALL-conformations which were set up to have no overlap, defined as more than 80% aligned sequence identity, with the training and validation sets.

As in the previous section, we evaluated ITsFlexible on crystal structures. However, as structural data is only available for a subset of known antigen receptor sequences (Olsen et al., 2022; Raybould et al., 2024), the model was additionally tested on predicted structural models (predicted with ImmuneBuilder (Abanades et al., 2023)). We compared model performance to the biophysical baselines and three zero-shot flexibility prediction workflows implemented based on the structure prediction tools AF2 (Jumper et al., 2021) and ABodyBuilder2 (ABB2) (Abanades et al., 2023). Two of the workflows made use of residue level confidence scores returned by these methods (pLDDT for AF2 and predicted error for ABB2) which have previously been described to provide a good indicator of disordered protein regions (Jumper et al., 2021). More specifically, CDR flexibility was classified from the mean AF2 pLDDT and the ABB2 root mean square predicted error (RMSPE) of loop residues. The third workflow predicts flexibility based on the diversity in CDRs modelled by multiple AF2 runs with subsampled MSAs (see methods for details).

ITsFlexible proved to be highly predictive of antibody and TCR CDR3 flexibility (Table 3). The method outperforms the biophysical baselines across all four CDR test sets. Similar performance is achieved for predictions from crystal structures and structural models.

**Table 3:**
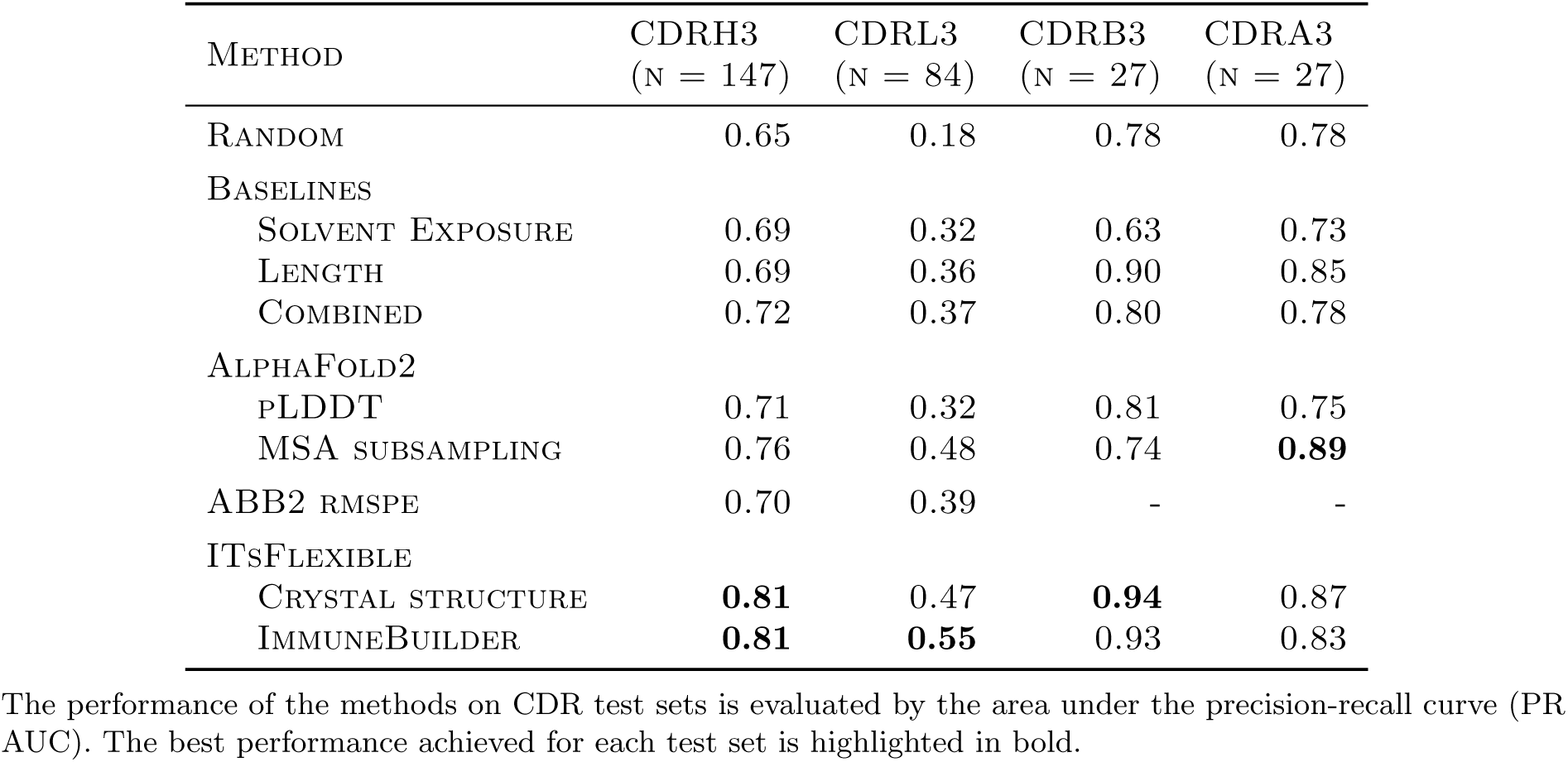
CDR test sets performance.

ITsFlexible also outperforms the AF2 and ABB2-based workflows (Table 3). The MSA subsampling approach surpasses ITsFlexible by a narrow margin on the CDRA3 set. However, ITsFlexible substantially outperforms all other methods on the CDRH3s, the largest and therefore most representative set, and is the only method that consistently achieves high predictive accuracy across all four test sets. A more detailed analysis of the ABB2 confidence score revealed that RMSPE is potentially more indicative of the number of times a certain antibody occurs in the training data than flexibility (see SI for details). Examples of CDRH3s predicted to be flexible and rigid are shown in Figure 2.

**Figure 2:**
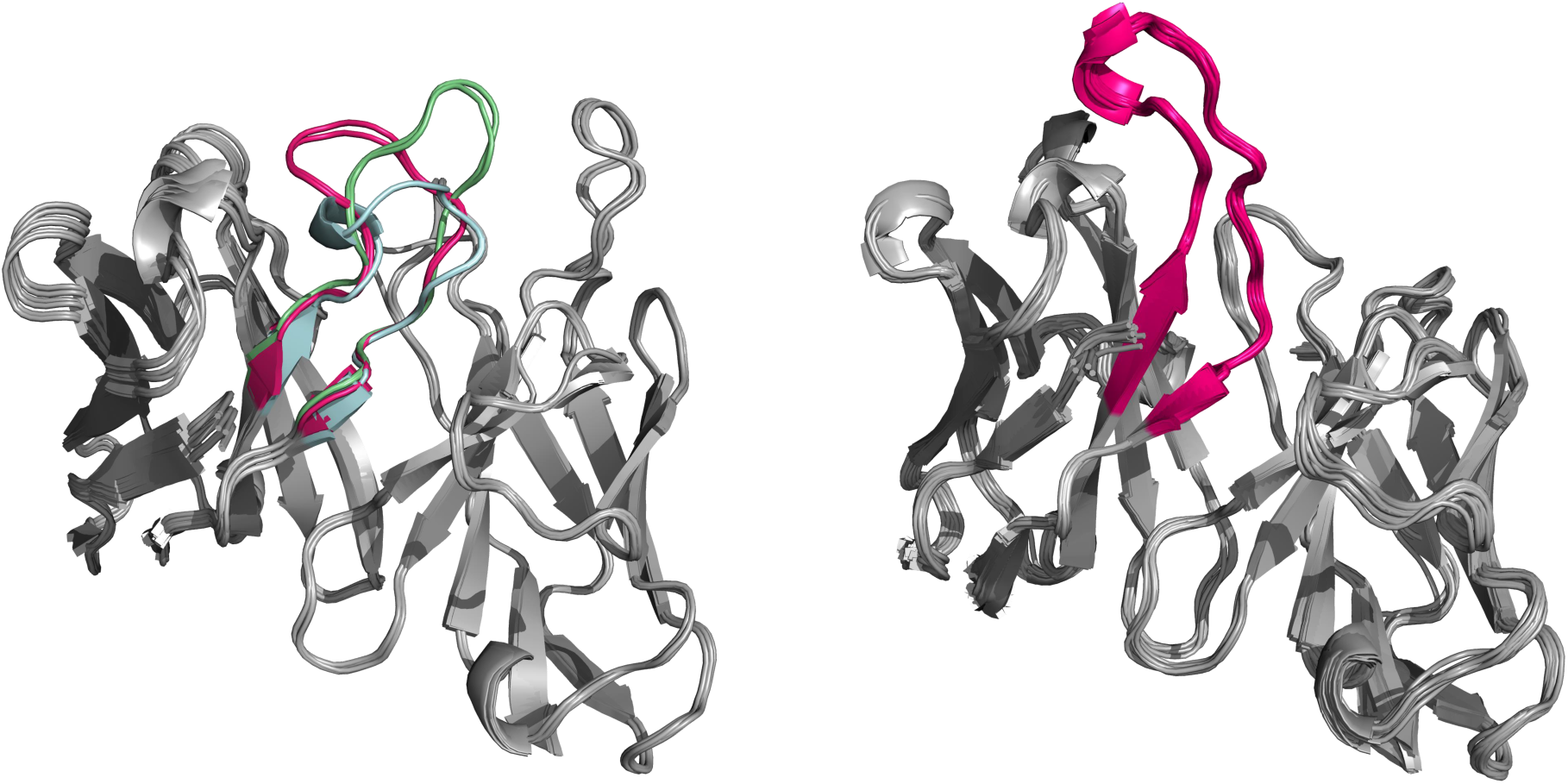
Example of antibodies predicted to be flexible and rigid by ITsFlexible. Left: Overlay of 6 structures of the same antibody Fv with CDRH3 predicted to be flexible. Crystal structures indicated that the CDRH3 (highlighted in colour) adopts three different conformations (red, blue, green). Right: Overlay of 22 structures of the same antibody Fv with CDRH3 predicted to be rigid. The CDRH3 (red) occupies the same conformations in all 22 structures.

**Figure 3:**
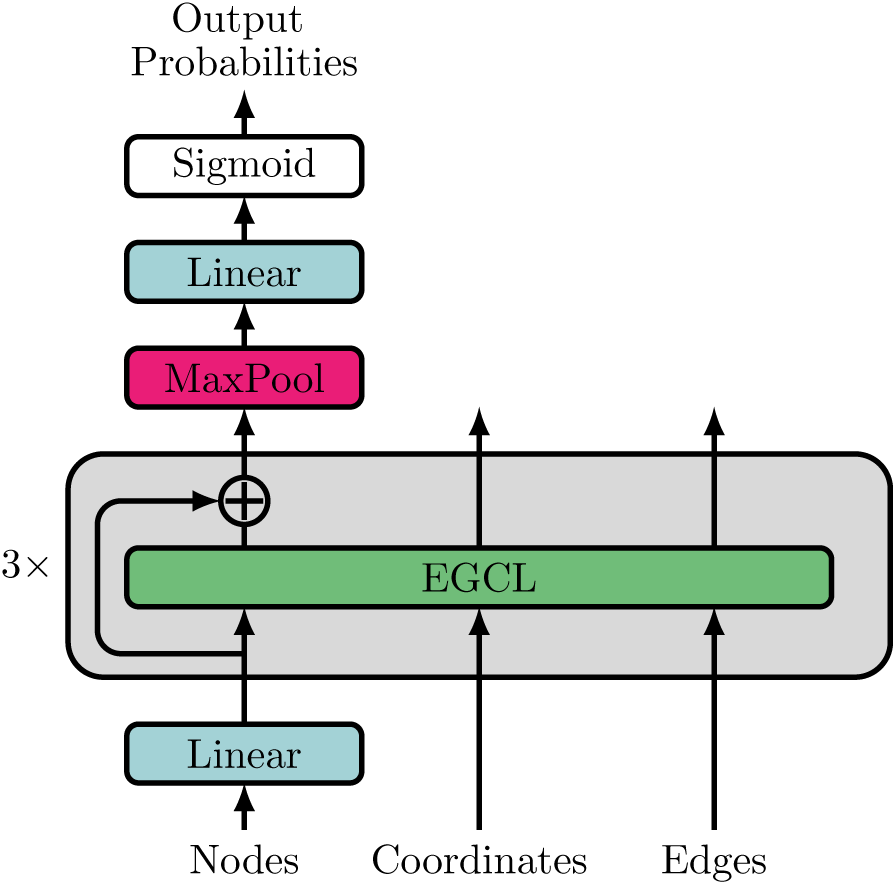
ITsFlexible model architecture. ITsFlexible is a GNN that consists of 3 E(n)equivariant graph convolutional layers (EGCLs). The final node features are max pooled and a classification head with linear layer and sigmoid activation is applied.

ITsFlexible was also evaluated with the most stringent train-test split. Specifically, CDRs in the ITsFlexible training set were filtered by 80% sequence identity to the test sets, compared to the 100% sequence similarity in the ABB2 training set, while the AF2 training set may even contain overlaps with the test sets.

Throughout the analysis in this manuscript we made the following assumptions to determine CDR flexibility. CDR3s are defined as IMGT residues 107-116 (see methods and Figure S3 for details) and structural diversity is calculated as the loop RMSD after alignment on the loop residues. We performed additional analysis on datasets with CDR3s defined by their exact secondary structure and flexibility calculated by alignment on the Fv and found that ITsFlexible’s high predictive accuracy remains qualitatively similar (see SI for details).

### 2.4 ITsFlexible matches CDR flexibility in molecular dynamics simulations

While crystallographic data can reveal conformational states of CDRs, it does not directly measure flexibility, and there always remains the possibility that additional conformational states exist but have not been captured. A limitation of the ALL-conformation dataset, therefore, is the lower confidence of ‘rigid’ compared to ‘flexible’ labels. Although we introduced requirements to address this limitation (see methods) the true extent of flexibility may well be underestimated. We also used MD simulations to classify CDR flexibility by simulating a set of 19 antibodies and labelling the flexibility of CDR3s (Table S2). As expected, we observed a higher proportion of flexible CDR3s in MD (84% for CDRH3, 37% for CDRL3) compared to the crystal structure data (64% for CDRH3, 18% for CDRL3). ITsFlexible achieved near perfect separation for CDRH3s, with slightly worse performance for CDRL3s (Table 4), suggesting that ITsFlexible also detects signals promoting flexibility in physics-based molecular simulations.

**Table 4:**
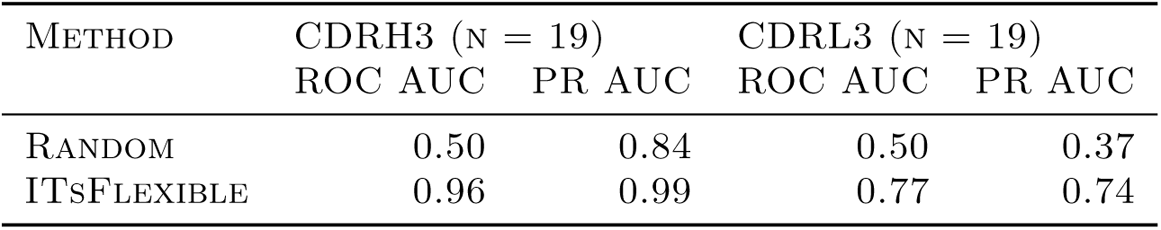
ITsFlexible performance on the MD test set.

### 2.5 Cryo-EM experiments confirm predicted flexibility

We chose three antibodies as challenging test cases for experimental validation using cryo-EM. Antibodies in the Patent and Literature Antibody Database (PLAbDab) (Abanades et al., 2024) against the influenza H1N1 hemagglutinin with ITsFlexible score below 0.1 (rigid examples) or above 0.5 (flexible examples) were considered. Three antibodies with low sequence identity to any examples in the training set and loop lengths opposing trends observed in ALL-conformations (long loop for rigid case, shorter loops for flexible cases) were selected (Table S13). These were imaged in complex with the antigen and the heterogeneity in the density maps, indicative of conformational variety (Punjani & Fleet, 2023), was analysed.

Antibody 1 is as an example of a long CDRH3 (length 19, longer than 86% of CDRH3s in ALL-conformation) predicted to be rigid with high confidence (ITsFlexible score of 0.02). The cryo-EM data revealed that the majority of good particles selected from 2D classification adopt one homogenous 3D class, which was used to build a high-resolution consensus structure (Figure S7). In agreement with model prediction this data shows that the CDR occupies a single conformation.

Antibody 2 is an example with a slightly shorter CDRH3 loop (length 16) predicted to be flexible with high confidence (ITsFlexible score of 0.76). Our initial data processing showed high flexibility in the binding interface of the antibody and the antigen albeit at low resolution. With further 3D classification and data collection, we were able to obtain a high resolution structure of one of the binding states (Figure S7). The resolution of the data did not allow us to build models of distinct conformational states which makes it hard to localize the flexibility to the CDRH3 loop. However, the density maps show clear conformational heterogeneity at the binding interface and induced by the antibody (Supplementary Movie 1 & 2). Flexibility of CDRH3 is a likely cause for this observation.

Antibody 3 is a further example of a CDRH3 predicted to be flexible. The CDRH3 is even shorter than in antibody 2 (length 13) and the ITsFlexible score of 0.60 suggests a lower prediction confidence. The cryo-EM data disagreed with the prediction and revealed no heterogeneity (Figure S7).

A potential limitation of the experimental setup is that antibodies were imaged in complex with the antigen. This choice was mad as the minimum particle size required for cryo-EM complicates the study of free antibodies. Binding can rigidify residues as additional molecular interactions with the antigen impose extra constraints (Mikolajek et al., 2022) and may prevent CDRs from accessing parts of the conformational space. ITsFlexible is trained on conformations of loop motifs irrespective of whether they interact with a binding partner and predictions are made from structures of unbound CDRs, we therefore expect predictions to better match the unrestricted conformational space more likely to be observed in the unbound antibody. This limitation could be a reason for the absence of flexibility observed in experiments for antibody 3. Accordingly, an argument could also be made that the unbound state of antibody 1 is more flexible than observed. The data, nevertheless, provide clear evidence for reduced flexibility of antibody 1 compared to antibody 2 when imaged under the same conditions, which matches our predictions.

Cryo-EM experiments of two challenging test cases provide further support that ITsFlexible has learned patterns associated with real CDR3 flexibility of antibodies in solution.

## 3 Discussion

Conformational changes are fundamental to functional properties of many proteins (Wei et al., 2016; Teilum et al., 2009). In antibodies and TCRs flexibility, especially of the highly important CDR3s, impacts key properties. CDR flexibility has been described to reduce affinity (Mikolajek et al., 2022; Schmidt et al., 2013) and promote polyspecificity (Guthmiller et al., 2020; James et al., 2003), both of which are generally undesired when engineering antigen receptors for use as therapeutics (Sormanni et al., 2018). A method predicting antigen receptor flexibility would, therefore, allow both an enhanced investigation of antibody function and the potential to tune desired therapeutic properties. However, current structure prediction tools, which excel at single structure prediction, are unable to model multiple conformational states of proteins and even more so antigen receptor CDRs reliably (Riccabona et al., 2024; Jing et al., 2024). One factor that has limited the progress in conformation prediction is the absence of large datasets necessary to train and evaluate such machine learning models.

Here, we focused on the computational prediction of antigen receptor CDR3 flexibility. To help overcome the lack of data we collected structural motifs with the same secondary structure pattern, defined as a loop connecting two consecutive antiparallel *β*-strands, across all proteins. Mining the PDB, we created ALL-conformations a dataset containing more than 1.2 million crystal structures of such loops with more than 100,000 unique sequences. Analysing the conformational flexibility in ALL-conformations we were able to label 20,000 loops with high confidence as either flexible, adopting multiple conformations, or rigid, occupying a single state. Using the ALL-conformations dataset we developed ITsFlexible, a method for the classification of CDR loops as either flexible or rigid. The model was trained on the subset of loop motifs from general proteins in the PDB and evaluated on its ability to predict CDR flexibility in an out-of-distribution setting. ITsFlexible achieved state-of-the-art performance on crystal structure test sets and generalised effectively to MD simulations data. In addition, we also showed that the model achieves similar performance when using inputs of structural models rather than crystal structures highlighting its applicability to antibodies without experimentally solved structures.

An ablation of ITsFlexible inputs indicated biophysical factors that influence the flexibility of CDRs. In line with previous MD studies (Guloglu & Deane, 2023), we identified the arrangement of residues in the structural context of CDRs as a key factor driving flexibility.

Furthermore, our analysis also showed that uncertainty scores of protein structure prediction tools and diversity in structural ensembles generated with established workflows are not reliable predictors of CDR flexibility. The AF2 pLDDT score has been described as a good indicator for disordered regions of proteins (Jumper et al., 2021), however, we found that it was not highly predictive of CDR flexibility. Similarly, the predicted error of the antibody specific ABB2 correlates more strongly with the number of times a particular sequence was present in the training set than flexibility. We also assessed the diversity of CDR ensembles produced by AF2 MSA subsampling as a predictor of flexibility. MSA subsampling is a popular workflow to model multiple conformational states of proteins (del Alamo et al., 2022; Stein & Mchaourab, 2022; Wayment-Steele et al., 2024; Sala et al., 2023b; Faezov & Dunbrack, 2023; Sala et al., 2023a; Saldaño et al., 2022). While this approach is more predictive than confidence scores, the workflow again does not capture CDR flexibility accurately.

We experimentally evaluated ITsFlexible’s predictive performance using three antibodies chosen as challenging test cases. The antibodies had low sequence identity to the training set and the loop length opposed trends observed in the data (long loop for rigid case, shorter loops for flexible cases). Using cryo-EM, we imaged the antibodies in complex with their antigen and analysed heterogeneity in the density maps (Zhong et al., 2021; Punjani & Fleet, 2023). For two of the three cases the experimental evidence confirmed our predictions. Among the correctly predicted cases, one rigid and one flexible, the former was three residues longer further demonstrating that the model detects features beyond loop length associated with flexibility. The third antibody featured an even shorter CDRH3 predicted to be flexible although with lower confidence. For this case study, experimental results showed no evidence of conformational heterogeneity. Overall, cryo-EM experiments provide strong evidence that ITsFlexible is predictive of real CDR3 conformational dynamics observed in solution.

In this work, we present ALL-conformations (available on Zenodo: doi.org/10.5281/zenodo.15032263) and ITsFlexible (available on Github: github.com/oxpig/ITsFlexible), two important resources to investigate and predict the conformational flexibility of antigen receptor CDRs. ALL-conformations captures the full range of experimentally observed conformational diversity of loops between antiparallel *β*-strands and enables training and benchmarking of more robust conformation prediction workflows. ITsFlexible accurately predicts CDR conformational flexibility and offers the potential to screen antigen receptors against undesired therapeutic properties. The development of these two resources opens the door to tackling more complex tasks such as sampling structures of different conformational states in the future.

## 4 Methods

### 4.1 ALL-conformations dataset

CDR3s of proteins in the immunoglobulin superfamily share a common secondary structure, they are formed by a loop that connects two antiparallel *β*-strands (Figure S3). We created **A**ntibody **L**ike **L**oop-conformations (ALL-conformations), a dataset consisting of five subsets that captures all observed conformational states of antibody CDRH3s and CDRL3s, TCR CDRB3s and CDRA3s and loop motifs between antiparallel *β*-strands across all proteins in the PDB. To generate the datasets, we implemented a systematic approach to search protein structure databases (PDB (Berman, 2000), SAbDab (Dunbar et al., 2014), STCRDab (Leem et al., 2018)) for all solved structures of the five protein motifs.

#### 4.1.1 The PDB set

We mined all the protein structures deposited in the PDB before November 22nd 2023 (Berman, 2000) for loop motifs sitting between two adjacent antiparallel *β*-strands. We used the DSSP algorithm (Kabsch & Sander, 1983) to assign secondary structure to amino acid residues. We then identified all antiparallel *β*-strands labelled by the algorithm and extracted the regions (loops) in between the two strands. The obtained loop structures were quality filtered for those solved by X-ray crystallography with resolution under 3.5 Å and no unresolved loop residues. A maximum of three residues forming secondary structure elements of *β*-strand or *α*-helix within each loop were allowed.

#### 4.1.2 Antibody and TCR CDR3 sets

All antibody Fv structures were extracted from the Structural Antibody Database (SAbDab) (Schneider et al., 2022; Dunbar et al., 2014). Both standard Fvs and single chain Fvs were included. All TCR Fv structures were extracted from the Structural T-Cell Receptor Database (STCRDab) (Leem et al., 2018). Multiple copies of antibodies in the same PDB structure were extracted as CDRs can adopt distinct conformations. Furthermore, some structures contain residues with alternate atom coordinates. These states were separated and individually added to the dataset. Structures were filtered for those solved by X-ray crystallography with a resolution below 3.5 Å, presence of a complete Fab (both heavy and light chain present in antibody structures; alpha and beta or gamma and delta chains for TCRs) and no unresolved residues in any of the CDRs.

Here, we defined CDR3 loops as the IMGT numbered (Lefranc et al., 2003) residues 107-116 which differs from the standard definitions. This choice was made to ensure that the definition of a CDR3 is more consistent with the way loops are defined by their secondary structure in the PDB set. The standard definition of a CDR3 loop in the IMGT numbering scheme are residues 105-117. However, an analysis of all antibody structures in SAbDab (Dunbar et al., 2014) showed that the residues at start and end of IMGT definitions tend to be part of the *β*-strands on either side of the loop (Figure S3). Positions 107-116 correspond better with the residues that sit between the two *β*-strands and we, therefore, used this range to define CDR3s throughout this manuscript.

#### 4.1.3 Conformational flexibility

We grouped the structures in ALL-conformations into sets of loops we consider identical. For the PDB set we grouped structures by sequence identity of the loop and considered all structures within a group to depict the same loop irrespective of the sequence of the rest of the protein. Antibody and TCR structures were grouped by sequence identity of the entire Fv rather than simply the CDR3s. In this way, CDR3s are only considered the to be the same if the entire domain is identical.

For loops with multiple available structures we analysed the conformational flexibility by calculating the number of accessible conformations. We defined a conformation based on structural similarity of a loop, using the RMSD of C*α* atoms. We clustered sequence identical loops using an agglomerate clustering algorithm with complete linkage (Murtagh & Contreras, 2012) and 1.25 Å distance threshold. This enforces that any two structures within a conformation have a maximum RMSD of 1.25 Å. This clustering approach was chosen as it is known to provide a good functional clustering of CDR loops in antibodies (Spoendlin et al., 2023). The structural similarity of loops can be calculated in multiple ways (see SI for another choice).

Each loop was assigned a label of ‘flexible’, ‘rigid’ or ‘unknown’. Loops for which multiple conformations (clusters) were found were assigned a flexible label. Loops for which only a single conformation was observed were not automatically assigned a ‘rigid’ label. The absence of multiple observed conformations does not prove that a loop cannot adopt multiple conformations. It is possible that alternative states were not captured by the limited number of crystal structures available. The more structures that are available of a loop in which it adopts a single conformation the more confident we can be on the absence of flexibility. Therefore, a requirement was introduced that a loop needs to adopt a single conformation in at least five separate PDB files to be labelled as rigid. We chose the requirement of five separate PDB files instead of simply five occurrences as the same loop can occur several times in the same PDB file due to multiple copies of a protein within a crystal unit cell. Loops neither labelled as ‘flexible’ or ‘rigid’ were assigned to the ‘unknown’ group.

### 4.2 ITsFlexible

ITsFlexible is a binary classifier. It takes as input a structural representation of a CDR loop motif and classifies it as flexible, able to adopt multiple conformations, or rigid, occupies a single stable conformation. Model architecture and training procedure are outlined below and details provided in the SI. The model is available on Github (github.com/oxpig/ITsFlexible).

#### 4.2.1 Model architecture

The ITsFlexible model is a graph neural network (Figure 3) and consists of three equivariant graph convolutional layers (EGCLs) (Satorras et al., 2022). The layers take input of node features, coordinates and edge features. Node features are iteratively updated and the last layer of node embeddings are pooled. A linear layer with sigmoid activation function is applied for binary classification. The chosen model architecture makes predictions invariant to transformations of the group E(3) (translations, rotations, reflections), therefore orientations and absolute positions of the input protein structure can be ignored. Predictions are dependent only on relative residue distances.

#### 4.2.2 Model inputs

A loop and its structural context were encoded as a residue level graph. The context was provided by all residues within 10 Å of any loop residues. For simplicity in the PDB set only residues located on the same protein chain as the loop were selected as context. For antibodies and TCRs residues located on both immunoglobulin chains were included, as the both chains influence conformational flexibility (Guloglu & Deane, 2023). Node features were a 22-dimensional vector consisting of a one-hot encoding of amino acid type (1 class for each of the 20 amino acids plus an additional class for unknown residues) and a one-hot encoding (1 class) whether the residue is located in the loop or structural context. Non-standard amino acids closely related to a standard amino acid were encoded as such, others were encoded as unknown residues (see Table S1). The amino acid encoding was concatenated with a one-hot encoding of the residue being located in the loop or the context. Coordinates for each node were taken as the position of the C*α* atom of a residue. Nodes were locally connected with edges using a 10 Å distance threshold. Edge features are 9-dimensional providing a one-hot encoding of the presence of a covalent bond between two residues and a C*α* distance encoding. The distance encoding was produced by 8 Gaussian radial basis functions (RBFs) equally distributed between 0 Å and 10 Å.

#### 4.2.3 Training

ITsFlexible was trained on the ALL-conformations PDB set using a 70-15-15 training, validation and test split. For length matched loops a maximal sequence identity of 80% was allowed between the splits. Additionally, all loops with more than 80% aligned sequence identity to any loop (not restricted to matching loop length) in the ALL-conformations CDRH/L/A/B3 sets were removed from the training and validation sets. One structure per loop was sampled for the validation and test sets. The training set was augmented by random sampling of five structures per loop to ensure the stability of predictions to small changes in atom coordinates. ITsFlexible was trained with a binary cross-entropy loss using the Adam optimiser (Kingma & Ba, 2014) with a learning rate of 2 *·* 10*^−^*^4^ and weight decay of 10*^−^*^6^. During training edges were dropped at random with a probability of 0.2. The validation area under the precision-recall curve (PR AUC) was monitored and training stopped when converged. Ten models were trained and the one with the best validation PR AUC selected.

### 4.3 Baseline models

A set of three baseline models were created to classify loop flexibility based on simple biophysical input features. The first baseline was formed by a logistic regression classifier fit to inputs of loop length. Longer loops contain more bonds around which they can rotate and are therefore expected to be more flexible in conformation than shorter ones (Guloglu & Deane, 2023; Marks et al., 2018). We found moderate correlation between loop length and flexibility in our datasets (Table S4). The second baseline was a logistic regression classifier fit to inputs of the solvent exposure of a loop. We approximated solvent exposure by the number of residues located within 10 Å radius around the loop. Loops with higher solvent exposure have less steric hindrance restricting conformational rearrangements and are expected to be more flexible (Guloglu & Deane, 2023). A final baseline model was formed by a logistic regression classifier fit to input of both length and solvent exposure. All baseline models were fit on the training split of the PDB set.

### 4.4 Alternative flexibility classifiers

Three alternative workflows were implemented to predict CDR flexibility. These were based on protocols designed to model protein conformational ensembles and confidence metrics of protein structure prediction tools.

#### 4.4.1 AlphaFold2 pLDDT

AF2 returns pLDDT scores for predicted structures which can be interpreted as a residue level confidence measure of the prediction. Antibody and TCR structures were predicted using the Colabfold implementation of AlphaFold-Multimer (Mirdita et al., 2022) with default parameters. The pLDDT score of the highest ranked model was extracted. The mean pLDDT of residues located in a CDR was used as input of a logistic regression model to classify flexibility.

#### 4.4.2 AlphaFold2 MSA subsampling

An AF2 MSA subsampling workflow, based on an established protocol (Riccabona et al., 2024; del Alamo et al., 2022), was used to predict structural ensembles. AlphaFold-Multimer (Evans et al., 2021) was run using the Colabfold (Mirdita et al., 2022) implementation to model antibody and TCR structures. We set the maximum MSA depth to 64, extra sequences to 128 and the number of recycles to 1 (default 3 in AF2). Low values of all three parameters increase the diversity of predicted structures and improves sampling of alternative conformations. Setting parameters too low can lead to the generation of unfolded structures (Riccabona et al., 2024). We chose these parameter values as they are within ranges described in previous studies (Riccabona et al., 2024; del Alamo et al., 2022) and they led to maximal diversity while limiting the occurrence of unfolded structures for the antibodies in our test set. Colabfold was run with 8 seeds resulting in 40 models (5 models are produced per seed) for each protein. The MSA subsampling protocol took approximately 3 minutes on a Nvidia A100 GPU. Owing to the computational cost, MSA subsampling was only performed for the smaller CDR sets and not the full PDB set.

We used the magnitude of structural diversity observed across the 40 predicted structures as a zero-shot classifier of CDR flexibility. Structural diversity was calculated by the C*α* RMSD of the CDR loop residues. The mean structural diversity was used as input of a logistic regression model to classify flexibility.

#### 4.4.3 ABB2 RMSPE

The ABB2 predicted error, a residue level confidence metric (Abanades et al., 2023), was used to classify flexibility of antibody CDRs. Antibody structures were predicted using a retrained version of ABB2. There is a large overlap of antibodies in the CDR test set used here for evaluation of flexibility prediction and the training set of ABB2. To avoid data leakage we retrained ABB2 removing all antibodies with 100% CDRH3 or CDRL3 sequence identity to any CDR in the test sets. This reduced the training set to 4469 antibodies compared to the 5669 in the original training set. This version of ABB2 retains good accuracy at antibody structure prediction evaluated on a benchmark (Table S11). For details on the ABB2 retraining as well as an evaluation of the original ABB2 for flexibility prediction see SI.

The predicted error of modelled antibodies was extracted. The root mean square predicted error (RMSPE) of CDR residues was used as input of a logistic regression model to classify flexibility.

### 4.5 Molecular dynamics simulations

Molecular dynamic simulations were performed according to the protocol described in FerńandezQuintero et al., 2019. The 19 investigated antibodies (Table S2) were prepared at pH 7.4 in MOE (ULC, 2024) using the Protonate3D tool (Labute, 2009). We capped C-termini using N-methyl groups and the structures were soaked in cubic water boxes of TIP3P water molecules with a minimum wall distance of 12 Å to the protein (Jorgensen et al., 1983; Gapsys & De Groot, 2020; Fischer et al., 2024). To neutralize the charges, the uniform background charge was applied, which is required to compute long-range electrostatic interactions (Sabri Dashti et al., 2012). To characterize different CDR loop conformational rearrangements and to broaden exploration of the conformational space, we employed well-tempered metadynamics implemented in GROMACS utilizing the PLUMED 2 software (Barducci et al., 2008; Domene et al., 2015; Tribello et al., 2014; The PLUMED consortium, 2019; Bonomi et al., 2009; Bauer et al., 2022). As collective variables we used a linear combination of the sine and cosine functions of the *ψ* torsion angles of the CDR loops (CDR-L3/H3 or all CDR loops). For our simulations, we employed a Gaussian height of 10 kJ/mol, with Gaussian deposition occurring every 5000 steps and a bias factor of 10. We performed 1000 ns of enhanced sampling simulations for each of the investigated systems. The obtained trajectories were aligned on the C*α*-atoms of the variable domains and clustered on the CDR loops using the average linkage hierarchical clustering algorithm with an RMSD cut-off criterion of 1.2 Å implemented in cpptraj, resulting in a large number of clusters(Roe & Cheatham, 2013). The respective cluster representatives were then equilibrated and used as starting structures for each 100 ns of classical molecular dynamics simulations using the AMBER 22 simulation package resulting in more than 10 µs of aggregated sampling time per system (Case et al., 2023).

MD simulations were performed in an NpT ensemble using pmemd.cuda (Salomon-Ferrer et al., 2013). Bonds involving hydrogen atoms were restrained by applying the SHAKE algorithm, allowing a time step of 2 fs. The Langevin thermostat was used to maintain the temperature during simulations at 300 K (Adelman & Doll, 1976; Doll et al., 1975) with a collision frequency of 2 ps^-1^ and a Monte Carlo barostat (Aqvist et al., 2004) with one volume change attempt per 100 steps.

Representative structures of the simulations were obtained by performing a hierarchical average linkage clustering implemented in cpptraj using an RMSD distance cut-off criterion of 2.5 Å for Fvs.

The number of conformations of simulated antibodies was determined as for the crystal structures datasets (see methods section: ALL-conformations datasets). The representative structures were clustered by C*α* RMSD of CDR3 loops using an agglomerative clustering algorithm with complete linkage (Murtagh & Contreras, 2012) and a 1.25 Å distance cutoff. The number of conformations observed for the CDR3s of each antibody are shown in Table S2. CDRH3s and CDRL3s were then assigned binary labels flexible, if multiple conformations were observed in the simulations, or rigid, in case of a single conformation.

The flexibility of simulated antibodies was predicted by ITsFlexible. A prediction was made for each representative structure of an antibody and the maximum prediction score used. We also tried using the mean prediction score, this however resulted in a more narrow distribution of scores across the dataset which is less optimal for classification tasks.

### 4.6 Cryo-EM experimental protocol and data analysis

The Patent and Literature Antibody Database (PLAbDab) (Abanades et al., 2024) was searched for antibodies that target influenza H1N1 hemagglutinin. Flexibility was classified with ITsFlexible from antibody models produced with ABB2 (Abanades et al., 2023). Antibodies with a predicted probability score lower than 0.1 were considered as rigid examples and antibodies with a score above 0.5 as flexible examples. Three antibodies (1 rigid, 2 flexible) with good developability properties predicted with TAP (Raybould et al., 2019) were selected for cryo-EM experiments.

#### 4.6.1 Sample preparation

3 HA (HA/California/07/2009) in complex with one Fab each (AEL31302/AEL31311 (antibody 1), AMB38310/AMB38599 (antibody 2) and AMB38442/AMB38568 (antibody 3) were prepared by mixing Fab with a 1:1 molar ratio of Fab:HA protomer and incubated for approximately 4 hours at 4°C. 0.25 µL of 4 mM CHAPSO detergent was added to the complex to aid in particle tumbling. The final concentration for the complexes of HA with each Fab were 1.14 mg/mL, 1.11 mg/mL and 1.02 mg/mL on the grid respectively. The samples were added to glow discharged 1.2/1.3 Au Ultrafoil 300 mesh grids and subjected to vitrification using a Vitrobot Mark IV system. The settings were as follows: 3 µL of sample, temperature inside the chamber was 4°C, humidity was 90%, blotting force was 1, wait time was 3 s. The sample was blotted off for 4.5 s and the grids were plunge-frozen into liquid-nitrogen-cooled liquid ethane.

#### 4.6.2 Data collection, processing, and model building

Cryo grids of the complexes were imaged at 190,000x nominal magnification using a Falcon 4i camera on a Glacios microscope at 200 kV. Automated image collection was performed using EPU from ThermoFisher. Micrographs for antibody 1 were collected untilted and micrographs for antibody 2 were collected with a 30° tilt. Images were aligned, dose-weighted, and Contrast Transfer Function (CTF)-corrected in the CryoSPARC Live^TM^ software platform, with automated image collection also performed using Smart EPU software (ThermoFisher). Data processing for all three datasets was carried out in CryoSPARC v4.5.3 (Punjani et al., 2017). Blob particle picking was performed on all micrographs with a minimum particle diameter of 100 Å and a maximum of 200 Å. Particles extracted at 480 pixels box size were used to perform 2D classification for antibody 1 and antibody 2, which were then used to generate a 3D reference model from ab-initio refinement, followed by heterogeneous refinement to obtain one good class that was further non-uniform (NU) heterogeneous refined. For antibody 3 we extracted particles at 512 pixels and Fourier cropped to 256 pixels box size to perform 2D classification, followed by ab-inito and heterogenous and NU refinements. Gold-Standard Fourier Shell Correlation (GSFSC) resolution was calculated to be 3.77 Å for antibody 1 and 3.91 Å for antibody 3. We docked the models into the cryo-EM density map in UCSF ChimeraX (Goddard et al., 2018). The structure model was built iteratively with COOT followed by real-space refinement in PHENIX package (Afonine et al., 2018). The Kabat numbering system (Kabat et al., 1979) was used for antibodies and H3 numbering scheme for HA.

For antibody 2 initially, no high-resolution map could be obtained as the antibody reveals a high degree of variability contributing to conformational changes in the head and stem of the HA respectively. The workflow to visualise the conformational changes in the complex is shown in Figure S1. We used the 243000 particles and the map resulting from the NU refine job downsized to 128 pixels as input for the 3D flex (Punjani & Fleet, 2023) data preparation job. For the 3D Mesh Prep job, a solvent mask was generated by low pass filtering the consensus map from 3D flex data prep job and segmented as shown in Figure S1 and a base number of 40 tetrahedral cells was used in combination with a minimum rigidity weight of 0.5. The resulting 3D flex mesh was used to run 3D flex train with a number of latent dimensions of 3. 3D Flex generator was used to produce a volume series of 41 frames using the consensus map showing the areas of high flexibility. The movie depicting the conformational flexibility based on the volume series was generated using UCSF ChimeraX (Goddard et al., 2018).

## Software and data availability

ITsFlexible method is publicly available on Github (github.com/oxpig/ITsFlexible). The ALL-conformations dataset and representative structures of MD simulated antibodies are released on Zenodo (doi.org/10.5281/zenodo.15032263). High-resolution cryo-EM structures of antibody 1 and antibody 2 were deposited in the PDB with codes 9N5Y and 9N5Z.

## Acknowledgements

This work was supported through research funding from the UK Engineering and Physical Sciences Research Council (EPSCR) [Grant number EP/S024093/1], Roche and the Royal Commission for the Exhibition of 1851 awarded to FS.

## Author contributions

FS and CD contributed to the conception and design of the study. FS created the crystal structure dataset, developed the ML models and performed model benchmarking. MF generated the MD data. MF, SR, HT, AG and JL performed the cryo-EM experiments and data analysis. FS and MF wrote the manuscript. FS, MF, SR, AB, WW, GG, and CD contributed to critical revision of the manuscript. All authors approved the submitted version.

