## Supplementary Information for "Predicting the conformational flexibility of antibody and T-cell receptor CDRs"

### S1 Supplementary methods

#### S1.1 ITsFlexible models

##### S1.1.1 ITsFlexible

ITsFlexible is an equivariant graph neural network (EGNN). The model takes a geometric graph as input that contains node features, coordinates and edge features and outputs a probability score that indicates the probability of a data point belonging to the positive class (details in Algorithm 1). Initially, node features are embedded into a 128 dimensional vector. Node embeddings are then updated through 3 E(n)-equivariant graph convolutional layers (EGCLs) (Satorras et al., 2022):

$$
\begin{aligned}
m_{ij} &= \phi_m(h_i^l, h_j^l, \|x_i^l - x_j^l\|^2, a_{ij}) \\
x_i^{l+1} &= x_i^l + C \sum_{j \neq i} (x_i^l - x_j^l)\ \phi_x(m_{ij}) \\
m_i &= \sum_{j \neq i} m_{ij} \\
h_i^{l+1} &= \phi_h(h_i^l, m_i)
\end{aligned}
$$

where $h^l$ are the node embeddings, $x^l$ are the coordinate and $a_{ij}$ the edge features. The last layer of node embeddings are maxpooled and a linear layer with sigmoid activation function is applied for binary classification. Coordinate updates in the EGCL are equivariant with respect to the group E(3) and node embedding updates are invariant. Classification of ITsFlexible is therefore invariant to translations and rotations of the input structure.

**Algorithm 1** ITsFlexible

**Require:** Node features $n \in \mathbb{R}^{N_{nodes} \times 22}$, node coordinates $x \in \mathbb{R}^{N_{nodes} \times 3}$, edges $a \in \mathbb{R}^{N_{edges} \times 9}$

```
def ITsFlexible(n, x, a)
    h_n^0 ← Linear(n_n)                          h_n ∈ R^128, n ∈ R ∩ [0, N_nodes]
    for l = 1 to 3
        a^l = DropoutEdge(a)
        h^l, x^l = EGCL(h^{l-1}, x^{l-1}, a^l)
        h^l = h^l + h^{l-1}
    end for
    h^4 = MaxPool(h^3)                           h^4 ∈ R^128
    o = Sigmoid(Linear(h^4))                     o ∈ R ∩ [0, 1]
    return o
```

##### S1.1.2 ITsFlexible-Loop

ITsFlexible-Loop is an EGNN very similar in architecture and training to ITsFlexible. The main difference is the model input. The ITsFlexible-Loop input is a graph encoding of only loop residues, the structural context is not provided. The coordinates and edge features of the graph are generated as described in the methods section in the main of this manuscript. As all nodes are located in the loop, node features are reduced to a 21-dimensional one-hot encoding of amino acid type. Hyperparameters of ITsFlexible-Loop were optimised independently. Node embeddings are 64-dimensional, the learning rate was set to $1 \cdot 10^{-3}$ and dropout to 0.1. There remaining hyperparameters are the same as for the ITsFlexible model.

##### S1.1.3 ITsFlexible-Sequence

ITsFlexible-Sequence is a CNN-based model which classifies loops from inputs of a sequence representation. Algorithm 2 shows details of the model. The model input is a one-hot encoding of the amino acid sequence. Each residue is encoded as 21 dimensional vector (1 class for each of the 20 amino acids plus an additional class for unknown residues). Non-standard amino acids closely related to a standard amino acid are encoded as such, others are encoded as unknown residues (Table S1). Encodings are padded with zeros to a standard length of 51 residues. This enforces a length limit of 51 residues for input sequences, which corresponds to the longest loop observed in the PDB (Berman, 2000). A sequence embedding is produced with two 1D-convolutional blocks with 256 channels. The final embedding is flattened to a vector of dimensions $51 \times 256$. Two linear layers followed by a sigmoid activation function are used for binary classification from the sequence embedding.

ItsFlexible-Sequence was trained on the PDB set with identical splits as ITsFlexible (see methods). ITsFlexible-Sequence was trained with a binary cross-entropy loss using the Adam optimiser (Kingma & Ba, 2014) with a learning rate of $2 \cdot 10^{-4}$. During training dropout of 0.2 was used in the convolutional layers and dropout of 0.05 in the linear layers. The validation area under the precision-recall curve (PR-AUC) was monitored and training stopped when converged. Ten models were trained and the one with the best validation PR-AUC selected.

**Algorithm 2** ITsFlexible-Sequence

**Require:** Loop sequence encoding $s \in \mathbb{R}^{N_{residues} \times 21}$

**def** ITsFlexibleSequence($s$)
  $i = \text{ZeroPad}(s)$  $i \in \mathbb{R}^{51 \times 21}$
  $h^0_n \leftarrow i_n$  $n \in \mathbb{R} \cap [0, 51]$
  **for** $l = 1$ **to** 2
    $h^l \leftarrow \text{ReLu}(\text{Conv1d}(h^{l-1}))$  $h^l \in \mathbb{R}^{51 \times 256}$
    $h^l = \text{Dropout}(h^l)$
    $h^l = \text{MaxPool1d}(h^l)$
  **end for**
  $h^3 \leftarrow \text{Flatten}(h^2)$  $h^3 \in \mathbb{R}^{(51 \times 256)}$
  $h^4 \leftarrow \text{Dropout}(\text{ReLu}(\text{Linear}(h^3)))$  $h^4 \in \mathbb{R}^{512}$
  $o \leftarrow \text{Sigmoid}(\text{Linear}(h^4))$  $o \in \mathbb{R} \cap [0, 1]$
  **return** $o$

#### S1.2 Amino acid encoding

When encoding ITsFlexible inputs, non-standard amino acids were represented either as a unknown residue or as the closest related standard amino acid. Non-standard amino acids that occur frequently in the test sets and are closely related to one of the 20 standard amino acids were encoded as such. Table S1 shows a map of the selected non-standard amino acids and their encoding.

Table S1: Encoding of non-standard amino acids

| Non-standard amino acid | Encoded as |
|---|---|
| Beta-L-aspartic acid (IAS) | Aspartic acid (ASP) |
| S-hydroxycystein (CSO) | Cystein (CYS) |
| 4-methyl-histidine (HIC) | Histidine (HIS) |
| N-dimethyl-lysine (MLY) | Lysine (LYS) |
| Selenomethionine (MSE) | Methionine (MET) |
| Phosphoserine (SEP) | Serine (SER) |
| Topaquinone (TPQ) | Tyrosine (TYR) |
| 3-amino-6-hydroxy-tyrosine (TYQ) | Tyrosine (TYR) |

#### S1.3 Molecular dynamics dataset

19 antibodies were simulated with our molecular dynamics protocol described in the methods section of this manuscript. The number of conformations observed for CDRH3s and CDRL3s is shown in Table S2.

Table S2: Number of conformations observed in molecular dynamics simulations

| Antibody | N CDRH3 conformations | N CDRL3 conformations |
|---|---|---|
| 1BFO | 3 | 1 |
| 1JPS | 3 | 1 |
| 1MLB D44 | 3 | 2 |
| 2Q76 D44 | 1 | 1 |
| 2VXT | 1 | 2 |
| 2Y06 | 3 | 1 |
| 2Y07 | 3 | 1 |
| 2Y36 | 3 | 1 |
| 3EOA | 5 | 1 |
| 3G6D | 3 | 4 |
| 3HI6 | 4 | 2 |
| 3RVW | 6 | 2 |
| 3V6F | 9 | 6 |
| 4KMT | 2 | 1 |
| 5I15 | 6 | 1 |
| 5I18 | 9 | 1 |
| 5I1A | 3 | 1 |
| 7G12 | 3 | 2 |
| 7G12 mature | 1 | 1 |

#### S1.4 Cryo-EM

Table S3: Cryo-EM data collection, refinement and validation statistics

| | antibody 1 - 9N5Y - AEL31302/AEL31311 | antibody 2 - 9N5Z - AMB38310/AMB38599 |
|---|---|---|
| **Data collection and processing** | | |
| Microscope | TFS Glacios | TFS Glacios |
| Voltage (keV) | 200 | 200 |
| Camera | TFS Falcon 4i | TFS Falcon 4i |
| Collection mode | Counting | Counting |
| Magnification | 190,000x | 190,000x |
| Pixel size at detector (Å) | 0.718 | 0.718 |
| Total electron exposure (e–/Å2) | 45 | 44.84 |
| Number of EER frames | 40 | 40 |
| Defocus range ($\mu$ m) | -0.8 to -1.6 | -0.8 to -1.6 |
| Automation software | EPU | EPU |
| Micrographs collected (no.) | 4,628 | 3,976 |
| Micrographs used (no.) | 4,457 | 3,295 |
| Initial particle images (no.) | 626,964 | 641,551 |
| Final particle images (no.) | 85,692 | 142,446 |
| Symmetry | C1 | C3 |
| Map resolution (masked/unmasked Å) | 3.7/3.8 | 3.0/3.1 |
| FSC threshold | 0.143 | 0.143 |
| **Refinement** | | |
| Initial model used (PDB code) | 7T3D | 7T3D |
| Refinement package | Phenix RSR | Phenix RSR |
| Model resolution (Å) | 3.77 | 3.16 |
| FSC threshold | 0.5 | 0.5 |
| EMRinger score | 3.08 | 3.32 |
| CC (mask) | 0.80 | 0.77 |
| Model composition | | |
| Non-hydrogen atoms | 13,754 | 13,702 |
| Protein residues | 1728 | 1724 |
| Ligands | 10 | 14 |
| R.m.s. deviations | | |
| Bond lengths (Å) | 0.005 | 0.002 |
| Bond angles (°) | 0.996 | 0.583 |
| Validation | | |
| MolProbity score | 1.39 | 1.61 |
| Clashscore | 3.37 | 12.41 |
| Poor rotamers (%) | 0.00 | 0.00 |
| Ramachandran plot | | |
| Favored (%) | 96.14 | 98.3 |
| Allowed (%) | 3.86 | 1.70 |
| Disallowed (%) | 0.00 | 0.00 |
| C$\beta$ outliers (%) | 0.00 | 0.00 |
| CaBLAM outliers (%) | 2.95 | 2.6 |

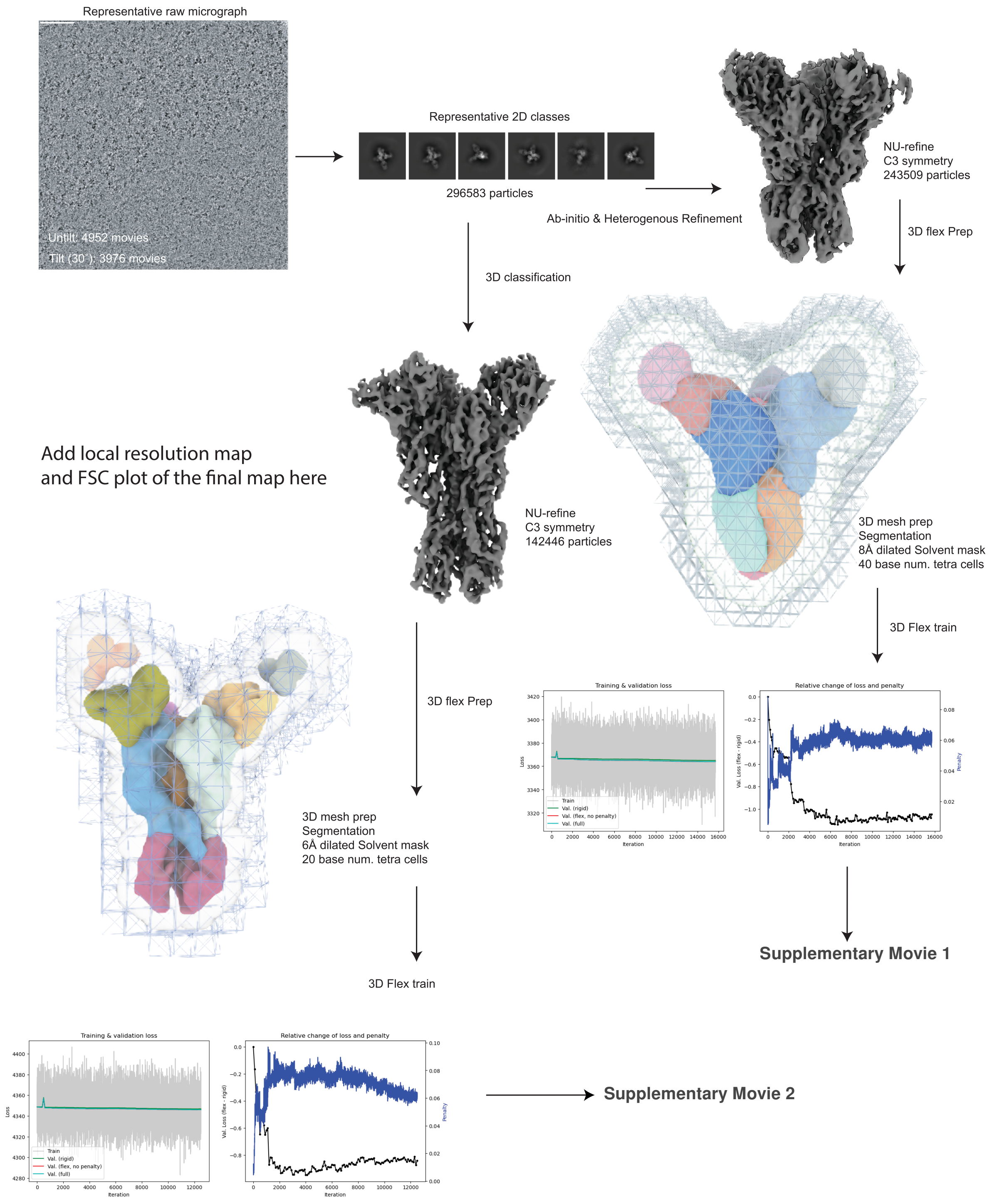

Figure S1: Workflow of the flexibility analysis in cryo-EM experiments. Figure detailing the 3D flex workflow described in the methods section of this manuscript.

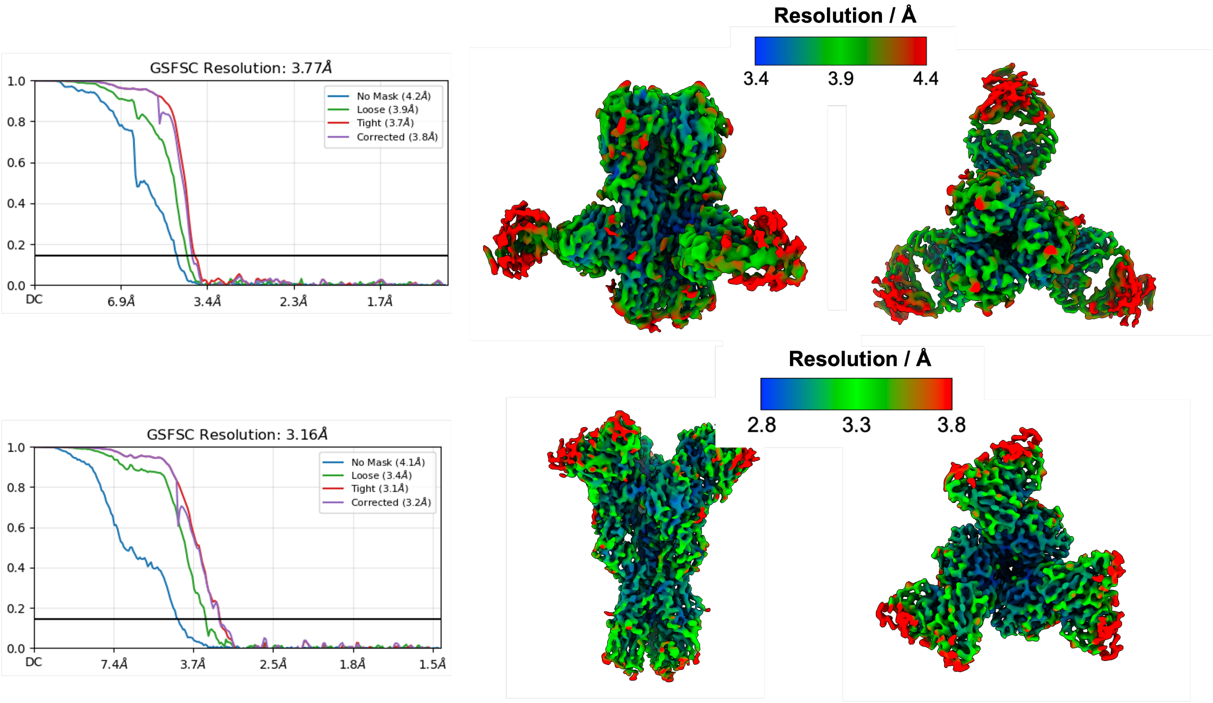

Figure S2: Local resolution maps for antibody 1 / 9N5Y (top) and antibody 2 / 9N5Z (bottom).

### S2 Supplementary results

#### S2.1 CDR3 secondary structure

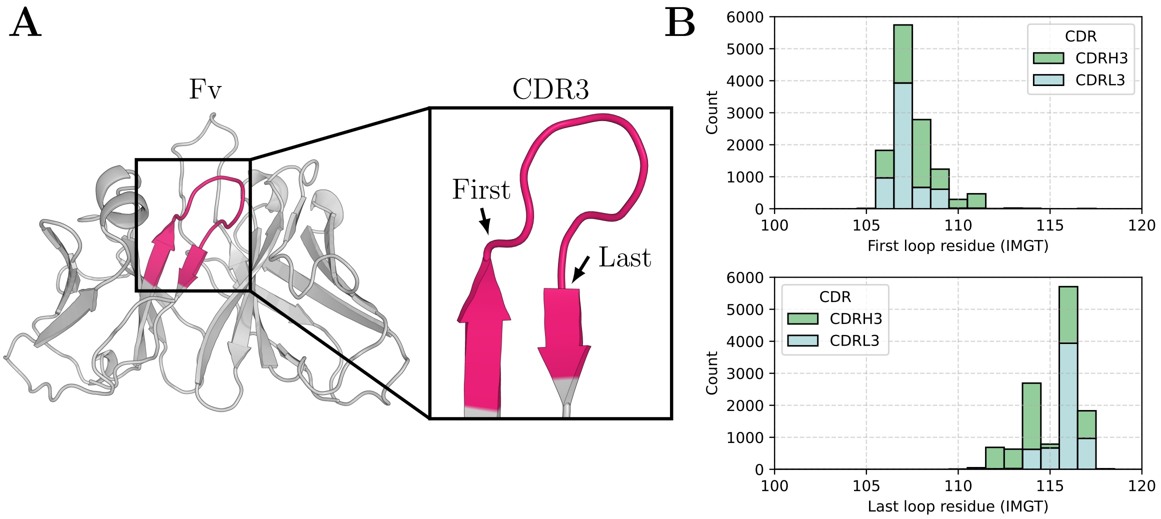

Figure S3: Antibody and TCRs CDR3 are loops bounded by two antiparallel $\beta$-strands. A) The variable domains (Fvs) adopt an immunoglobulin fold which consist of 9 antiparallel $\beta$-strands. CDR3s, defined by IMGT residues 105 - 117 (highlighted in pink), connect the two C-terminal strands of the Fv. The start (105) and end (117) residues are located on the strands and the majority of the CDR3 forms a loop motif. The first and last residue in the loop motif are highlighted. B) Histogram showing the distribution of antibody variable domain residues (IMGT numbered) that occur as the first and last residue of the loop motif. The majority of loops start at IMGT residue 107 and end at IMGT residue 116. Unless stated otherwise, we define a CDR3 as IMGT residue 107-116 throughout the paper.

#### S2.2 ALL-conformations

##### S2.2.1 ALL-conformations statistics

This section provides additional statistics for ALL-conformations. The distributions of loop length and the number of structures available for loops are shown in Figure S4 & S5. Correlation between flexibility and loop length is shown in Table S4. The number of loops labelled as 'flexible' and 'rigid' in each set is shown in Table S5

Table S4: Correlation of CDR length and flexibility

| Set | Original dataset (alignment on CDR) | Alternative dataset (alignment on Fv) |
|---|---|---|
| CDRH3 | 0.13 | 0.11 |
| CDRL3 | 0.34 | 0.21 |
| CDRB3 | 0.36 | 0.16 |
| CDRA3 | 0.12 | -0.03 |

R values of the point biserial correlation of the number of residues in the CDR and binary labels rigid (0) and flexible (1).

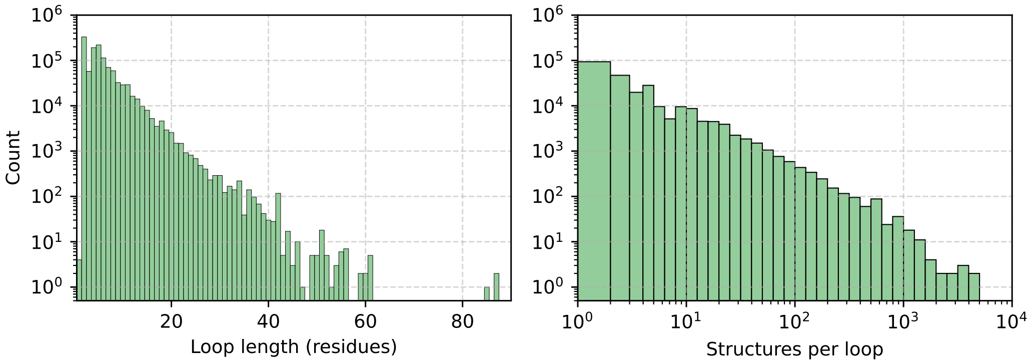

Figure S4: Length distribution and number of available structures for loops in the PDB set of ALL-conformations.

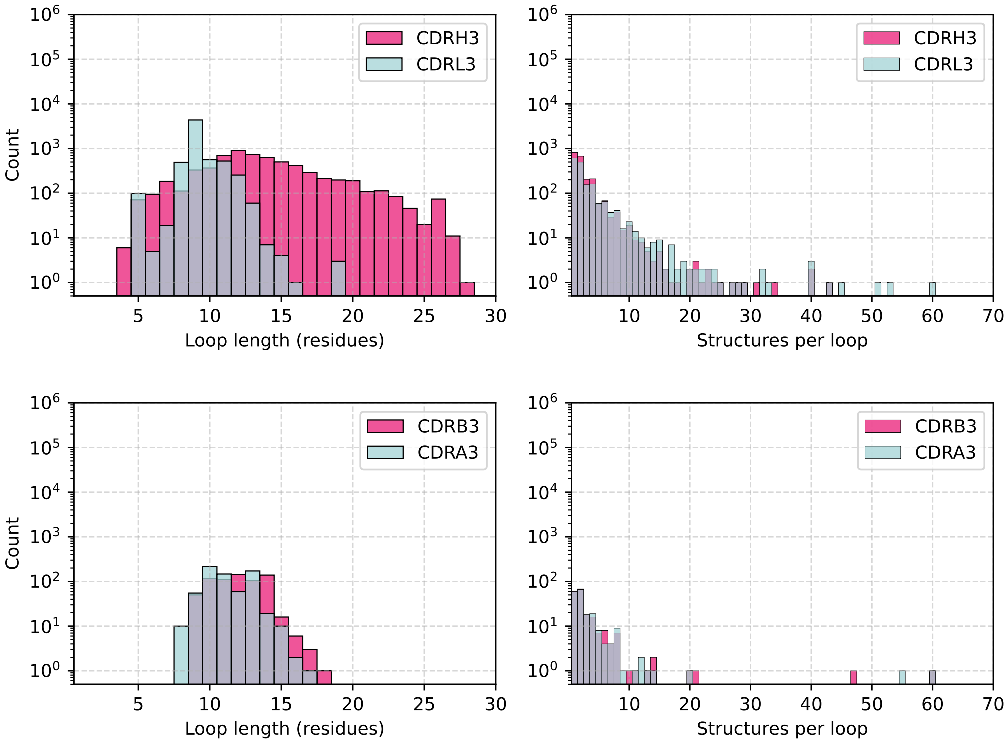

Figure S5: Length distribution and number of available structures for CDRs in ALL-conformations. top) antibodies CDRH3 and CDRL3 sets, bottom) TCRs CDRB3 and CDRA3 sets.

Table S5: Conformational flexibility in ALL-conformations

| Dataset | Total loops with determine flexibility | Multiple observed conformations - flexible | Single observed conformation - rigid |
|---|---|---|---|
| PDB set | 20,814 | 4289 | 16,525 |
| Antibodies | | | |
| CDRH3 | 152 | 97 | 55 |
| CDRL3 | 84 | 15 | 69 |
| TCRs | | | |
| CDRB3 | 27 | 21 | 6 |
| CDRA3 | 27 | 21 | 6 |

###### S2.2.2 Alternative definition of loop flexibility

In the main sections of this manuscript, we made several choices to determine the conformational flexibility of loops in ALL-conformations. Structures in the PDB set were initially grouped by sequence identity of the loop, whereas structures in the CDR3 sets were grouped by sequence identity of the Fv. Structural similarity was then calculated by the RMSD of loop residues after structural alignment on the loop.

In this section, a different approach to determine the conformational flexibility was explored. In the PDB set structures were grouped by sequence identity of the loop and two additional anchor residues on either of its sides. Structural similarity was then calculated by the RMSD of the loop after structural alignment on the anchor residues.

Antibody and TCR structures were grouped by Fv identity as in the original dataset. To calculate structural similarity, structures were aligned on the framework regions of the chain the CDR is located. We chose to align on the entire framework region rather than anchor residues (as for the PDB set) to reduce noise in the alignment. In certain cases the $\beta$-strand located on the C-terminal side of the CDR3 is not resolved well in crystal structures due to its proximity to the end of the chain. When choosing to align on anchor residues only the alignment can therefore show inaccuracies.

A conformation was defined, as described in the methods section, by clustering using an agglomerative clustering with complete linkage and 1.25 Å distance threshold. We recalculated the flexibility of loops in ALL-conformations using this alternative definition of flexibility. Summary statistics for are shown in Table S6.

Table S6: Conformational flexibility in ALL-conformations using alternative definitions

| Dataset | Total loops with determined flexibility | Multiple observed conformations - flexible | Single observed conformation - rigid |
|---|---|---|---|
| PDB set | 34,453 | 18,526 | 15,927 |
| Antibodies | | | |
| CDRH3 | 236 | 187 | 49 |
| CDRL3 | 127 | 66 | 61 |
| TCRs | | | |
| CDRB3 | 41 | 35 | 6 |
| CDRA3 | 36 | 31 | 5 |

##### S2.3 ITsFlexible performance with secondary structure-based CDR3 definitions

Throughout the analysis in the main sections of this manuscript we defined CDR3s as IMGT residues 107-116. This range was shown to align with the secondary structure of the loop for the majority of antibodies (Figure S3). Here, we assessed the impact of defining each CDR3 exactly by its secondary structure. We used the DSSP algorithm (Kabsch & Sander, 1983) to identify the exact numbering of the first residue after the immunoglobulin F-strand and the last residue before the G-strand and extracted the region in between as the CDR3. We refer to this definition as the DSSP CDR3.

CDRH3 and CDRL3 test sets were created with conformational flexibility recalculated over DSSP CDR3 definitions and ITsFlexible performance was evaluated (Table S7). Similar performance is achieved for the DSSP and our standard CDR3 definitions. Due to the simplicity of CDR3s being defined by the same IMGT numbers in all antibodies across the dataset, we chose to use the 107-116 definition throughout the manuscript.

Table S7: DSSP defined CDR test set performance.

| Method | CDRH3 | | CDRL3 | |
|---|---|---|---|---|
| | DSSP | IMGT 107-116 | DSSP | IMGT 107-116 |
| Random | 0.62 | 0.65 | 0.19 | 0.18 |
| Baselines | | | | |
| Solvent Exposure | 0.66 | 0.69 | 0.25 | 0.32 |
| Length | 0.64 | 0.69 | 0.29 | 0.36 |
| Combined | 0.73 | 0.72 | 0.38 | 0.37 |
| ITsFlexible | | | | |
| Crystal structure | **0.79** | **0.81** | **0.50** | **0.47** |

The performance of the methods on CDR test sets is evaluated by the area under the precision-recall curve (PR AUC). The best performance achieved for each test set is highlighted in bold.

##### S2.4 ITsFlexible performance with alternative definition of flexibility

ITsFlexible was retrained on the dataset with alternative definitions of flexibility as described in Section S2.2.2. This model was parametrised similar to the model described in the methods section. The context distance threshold was increased to 30 Å and the edge threshold distance to 20 Å to provide the model with information of a larger structural context.

Model performance was evaluated on the PDB (Figure S8) and the CDR test sets (Figure S9). The data splits were created based on sequence identity. The PDB test set did not contain any loops with more than 80% sequence identity across the loop and anchor residues to length matched loops in the training and validation sets. The CDR test sets did not contain any loops with more than 80% aligned sequence identity to any loop (not restricted to matching loop length) in the training and validation sets. Baselines and alternative workflows were performed as outlined in the methods. We made a small modification of the AF2 MSA subsampling to make the workflow more consistent with the here used definition of flexibility. Flexibility in the structural ensembles was calculated by loop RMSD after alignment on the framework regions of the corresponding chain. ITsFlexible achieves good preformance on all five datasets indicating the models ability to detect flexibility signals independent of choices made when calculating structural flexibility.

Table S8: PDB test set performance.

| Method | ROC AUC | PR AUC |
|---|---|---|
| Random | 0.50 | 0.55 |
| Baselines | | |
| Solvent exposure | 0.60 | 0.64 |
| Length | 0.64 | 0.64 |
| Combined | 0.74 | 0.75 |
| ITsFlexible | | |
| ITsFlexible-sequence | 0.70 | 0.71 |
| ITsFlexible-loop | 0.65 | 0.68 |
| ITsFlexible | **0.81** | **0.82** |

Table S9: CDR test set performance

| Method | CDRH3 (n = 236) | CDRL3 (n = 127) | CDRB3 (n = 41) | CDRA3 (n = 36) |
|---|---|---|---|---|
| Random | 0.79 | 0.52 | 0.85 | 0.86 |
| Baselines | | | | |
| Solvent Exposure | 0.85 | 0.67 | 0.82 | 0.84 |
| Length | 0.84 | 0.70 | **0.91** | 0.88 |
| Combined | 0.87 | 0.73 | 0.88 | 0.82 |
| AlphaFold2 | | | | |
| pLDDT | 0.82 | 0.66 | 0.88 | 0.80 |
| MSA subsampling | 0.87 | 0.56 | 0.87 | 0.89 |
| ABB2 rmspe | 0.76 | 0.70 | - | - |
| ITsFlexible | | | | |
| Crystal structure | **0.88** | **0.83** | 0.90 | **0.90** |
| ImmuneBuilder | **0.88** | 0.79 | 0.87 | 0.86 |

The performance of the methods on CDR test sets is evaluated by the area under the precision-recall curve (PR AUC). The best performance achieved for each test set is highlighted in bold.

#### S2.5 ABodyBuilder2

##### S2.5.1 ABB2 retraining

The original ABB2 (Abanades et al., 2023) was trained on a version of SAbDab (Schneider et al., 2022) downloaded in July 2021. As SAbDab was also used to create ALL-conformations and consequently the antibody CDR flexibility test sets, there is a substantial overlap with the ABB2 training set (Figure S10). To avoid data leakage, ABB2 was retrained using the original training protocol on a dataset with the ALL-conformations overlap removed. Antibodies with 100% sequence identity in CDRH3 or CDRL3 regions to a data point in the CDRH3 or CDRL3 flexibility test sets were removed from the training set. This reduced the number of training set antibodies from 5669 to 4469. The validation and test set were not changed. The accuracy of the retrained ABB2 is compared to the original ABB2 on a benchmark of 34 non-redundant antibody structures from the SAbDab (Table S11). The retrained ABB2 retained high accuracy at predicting antibody structures.

Table S10: Overlap of antibody CDR test sets and ABB2 training set

| | CDRH3 | CDRL3 |
|---|---|---|
| Total in test set | 147 | 84 |
| ABB2 training overlap* | 127 | 76 |

*Number of CDR3 sequences that also appear in the ABB2 training set.

Table S11: Comparison of ABB2 and retrained ABB2 on benchmark

| Method | CDRH1 | CDRH2 | CDRH3 | FwH | CDRL1 | CDRL2 | CDRL3 | FwL |
|---|---|---|---|---|---|---|---|---|
| ABB2 original | 0.86 | 0.74 | 2.61 | 0.74 | 0.52 | 0.38 | 0.87 | 0.56 |
| ABB2 retrained | 0.87 | 0.75 | 2.93 | 0.76 | 0.54 | 0.35 | 0.92 | 0.58 |

The mean RMSD (in Å) of the prediction to the crystal structure across the benchmark set is shown for the six CDRs and the framework regions according to IMGT definitions.

##### S2.5.2 Flexibility classification with ABB2

In this section, we show results of using the original ABB2 (trained by Abanades et al. (2023)) for flexibility classification and highlight that due to data leakage and dataset biases the performance is not representative. Flexibility classification based on RMSPEs obtained from structures predicted with the original ABB2 performs substantially better than using the retrained ABB2 (Table S12). This boost in performance is linked to biases in the CDR flexibility test sets. When labelling CDRs as rigid or flexible, we introduced a requirement that a rigid CDR needs to appear in at least five separate PDB structures (see methods). Therefore, CDRs labelled as rigid tend to be represented by more copies in the ABB2 training set as the ones labelled to be flexible (Figure S6).

We further observed a negative correlation between the number of times a CDR occurs in the ABB2 training set and its RMSPE (Figure S6). This finding can be explained by the way that ABB2 calculates the PE score. ABB2 predicts four structures for each antibody, the PE score is then calculated as the diversity between these structures (Abanades et al., 2023). The more often an antibody appears in the training set, the more it is weighted in the training loss. This makes it more likely that the antibody structure is predicted with high accuracy and reduces the diversity between the four structures. The correlation disappears for the retrained ABB2 which shows that the correlation is indeed an artifact of the training set.

To highlight the importance of this bias, we predicted flexibility simply based on the number of times an antibody appears in the original ABB2 training set (Table S12). This predictor, based on data bias only, performs closer to the original ABB2 than to the random baseline.

Table S12: Flexibility classification with ABB2

| Method | CDRH3 (n = 147) | CDRL3 (n = 84) |
|---|---|---|
| Random | 0.65 | 0.18 |
| ITsFlexible | 0.81 | 0.55 |
| ABB2 rmspe | | |
| Retained | 0.70 | 0.39 |
| Original | 0.81 | 0.62 |
| N structures in ABB2 training set* | 0.74 | 0.41 |

*Logistic regression model with input corresponding to the number of times a given CDR occurs in the ABB2 training.
The performance of the methods on CDR test sets is evaluated by the area under the precision-recall curve (PR AUC).

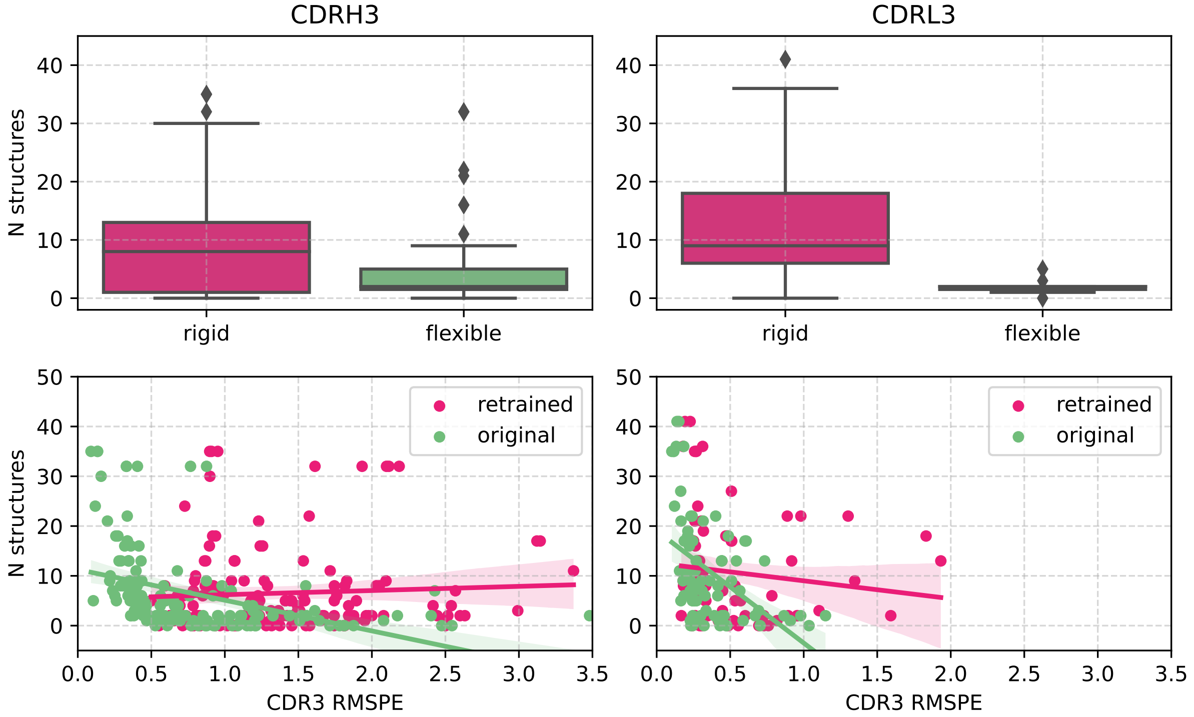

Figure S6: Biases in the antibody CDR3 flexibility classification test sets. top) Boxplot of the number of structures appearing in the original ABB2 training set for CDRs labelled as flexible and rigid. Rigid CDRs tend to be represented more often in the training set than flexible structures. bottom) The number of structures in the original ABB2 training set is plotted against the CDR3 RMSPE for the retrained and the original ABB2. A linear regression is fit to the data points. For the original ABB2 there is a moderate negative correlation (CDRH3: $R = -0.42$, CDRL3: $R = -0.47$) between the number of training set structures and the RMSPE of a CDR. For the retrained ABB2 there is no correlation (CDRH3: $R = 0.06$, CDRL3: $R = -0.12$).

#### S2.6 Case study antibodies and high-resolution cryo-EM structures

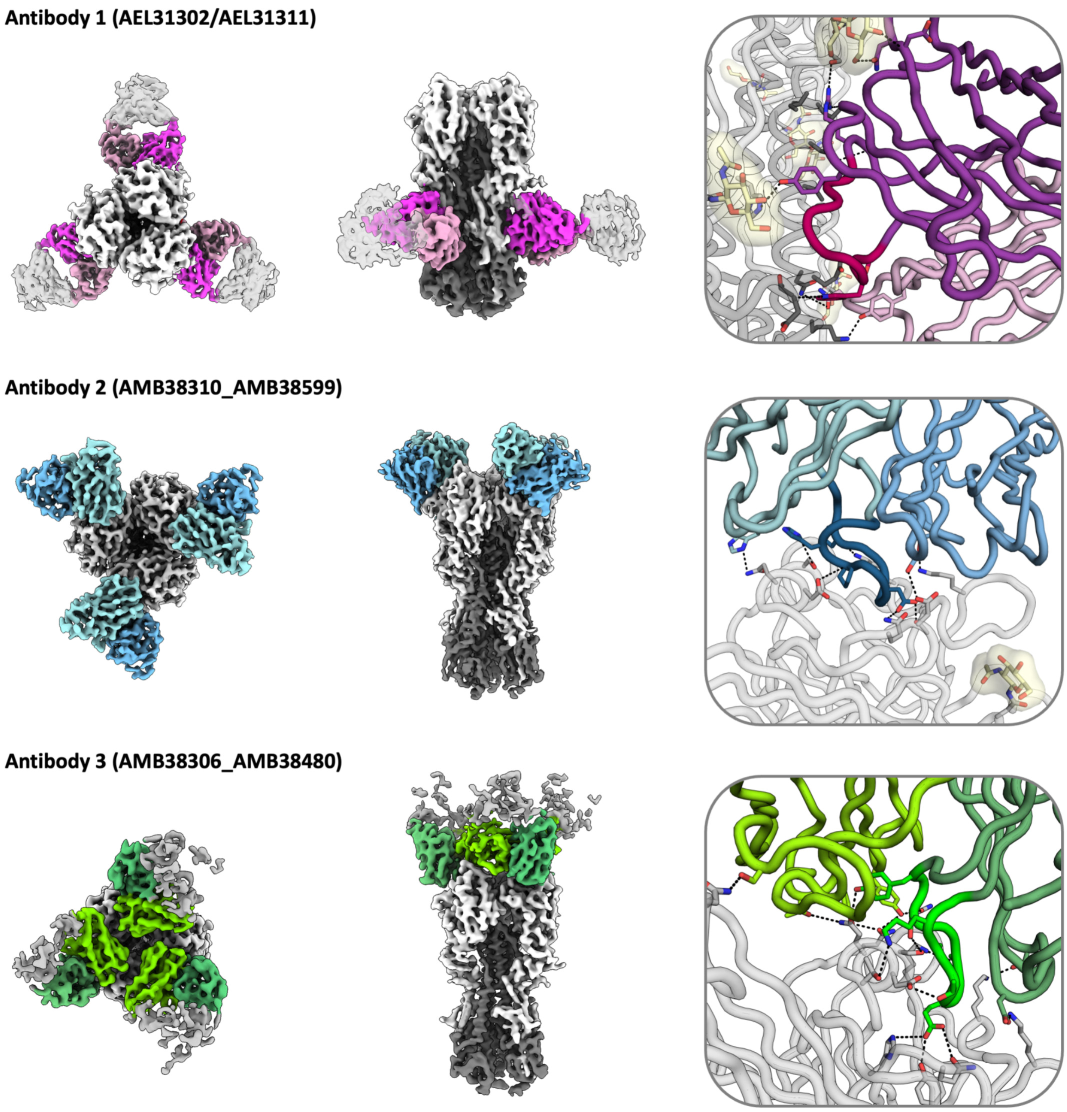

Figure S7: High-resolution structure of the three case study antibodies in complex with influenza H1N1 hemagglutinin. Antibody 1 and 2 were deposited in the PDB with codes 9N5Y and 9N5Z. (Left) Top and side view of the cryo-EM map of antibodies (colour, heavy chain dark, light chain bright) in complex with the antigen (grey). In each structures three symmetrically arranged copies of the antibody were captured. Antibody 1 binds to the hemagglutinin stem and antibodies 2 and 3 to the hemagglutinin head. (Right) Cartoon representation of antibody-antigen binding interface. CDRH3s are shown in the different shade of colouring and binding interactions are highlighted in stick representation.

Table S13: Antibodies selected for cryo-EM experiments

| ID | CDRH3 length | CDRH3 seq identity to train set | ITsFlexible score | Prediction | Cryo-EM |
|---|---|---|---|---|---|
| 1 - 9N5Y | 19 | 0.32 | 0.02 | rigid | rigid |
| 2 - 9N5Z | 16 | 0.44 | 0.76 | flexible | flexible |
| 3 | 13 | 0.46 | 0.60 | flexible | rigid |
